# Deregulation of epigenetic marks is correlated to differential exon usage of developmental genes

**DOI:** 10.1101/2020.12.17.423086

**Authors:** Hoang Thu Trang Do, Siba Shanak, Ahmad Barghash, Volkhard Helms

**Affiliations:** Saarland University, Center for Bioinformatics, 66041 Saarbruecken, Germany; Arab American University, Dept. for Biology and Biotechnology, Jenin, Palestine; German Jordanian University, School of Electrical Engineering and Information Technology, Amman, Jordan

## Abstract

Alternative exon usage is known to affect a large portion of genes in mammalian genomes. Importantly, different splice forms sometimes lead to distinctly different protein functions. We analyzed data from the Human Epigenome Atlas (version 9) whereby we connected the differential usage of exons in various developmental stages of human cells/tissues to differential epigenetic modifications at the exon level. In total, we analyzed 19 human tissues, adult cells, and cultured cells that mimic early developmental stages. We found that the differential occurrence of protein isoforms across developmental stages was often associated with changes in histone marks at exon boundary regions. Many of the genes that are differentially regulated at the exon level were found to be functionally associated with development and metabolism.

## INTRODUCTION

Alternative splicing (AS) or differential exon usage (DEU) was reported to occur in 90-95% of all human multi-exon genes [1, 2] and leads to a strong expansion of the eukaryotic proteome [3]. AS is an integral part of differentiation and developmental programs and contributes to cell lineage and tissue identity as reported by Wang et al. for nine different human tissues [4]. Based on the transcriptomes of 15 different human cell lines, the ENCODE project reported that up to 25 different transcripts can be produced from a single gene and up to 12 alternative transcripts may be expressed in a particular cell [5], respectively.

It is well established that AS is tightly associated with respective epigenetic chromatin modifications [6–9]. The role of chromatin in AS was first suggested by Adami and colleagues who found that two copies of the same adenovirus genome in the same nucleus gave rise to different spliced RNAs [10]. Another well-documented example where H3K36me3 influences AS of a mammalian transcript is the fibroblast growth factor receptor (FGFR2). FGFR2 was reported to accumulate histone modifications H3K36me3 and H3K4me1 along the alternatively spliced region in mesenchymal cells, where exon IIIc is included. In contrast, H3K27me3 and H3K4me3 were found to be enriched in epithelial cells, where exon IIIb is used [11]. In mesenchymal cells, H3K36me3 is recognized by the MRG15 protein that recruits the splicing factor PTB to the intronic splicing silencer element surrounding exon IIIB to repress its inclusion in these cells [11].

The relationship between DEU and differentiation / development has also been extensively addressed. Kalsotra and Cooper reviewed in 2011 the current understanding of the roles of AS in cell division, cell fate decisions, and in tissue maturation [12]. More recently, Baralle and Giudice reviewed the connection between AS and cell differentiation as well as with epigenetic landscapes, and the role of splicing processes in the brain, striated muscle, and other tissues and organs [13]. More focused studies addressed, for example, how the splicing regulators Esrp1 and Esrp2 direct an epithelial splicing program that is essential for mammalian development [14], and the role of splicing for neural development [15]. Although the pairwise connections between AS and epigenetic modifications and between AS and differentiation / development have each been characterized in detail, the circular connection between differential exon usage, epigenetic modifications and development received apparently relatively little attention so far. As mentioned, Baralle and Guidice summarized some work describing such interplay in neurological and brain development [13]. An interesting study from the Heller lab related the enrichment of histone post-translational modifications to AS regulation during tissue development in mice. They found, for example, that enrichment of histone modifications H3K36me3 and H3K4me1 in exon flanking regions was wired to skipped exon selection with strong evidence across all investigated embryonic tissues and developmental time points [16].

The exon-intron boundary was shown to be the most important region for epigenetically regulated AS. For example, Guan et al. reported strong association between epigenetic signals and cassette exon inclusion levels in both exon and flanking regions [17]. In another example, flanking areas annotated with exon skipping and alternative splice site selection events were found to be statistically enriched with DNA methylation, nucleosome occupancy and histone modification [18]. In a recent study based on ENCODE human data, Gerstein and co-workers used spatio-temporal epigenetic features extracted from exon flanks to model splicing regulation, and identified H3K36me3, H3K27me3, H3K4me1, H3K4me3, H3K9me3 and H3K27ac as highly influential features in the splicing regulatory model [19].

The authors of these previous studies, however, did not explicitly relate their findings to developmental stages and this is precisely the aim of this study. Based on data from the Roadmap Epigenomics Project for human tissues and adult and cultured stem cells, we aimed at correlating differential exon usage to epigenetic modifications of different histone marks at the exon boundaries. We found strong correlations for alternatively spliced genes and identified subsets of genes that are highly expressed in specific tissues. This then enabled us to examine splicing regulation patterns shared between the investigated cells. Furthermore, we were able to associate the co-occurrence of differential exon usage and histone modifications with functional annotations that, indeed, are often related to regulation of signaling and developmental processes. Besides a global analysis, we also present a detailed analysis of the two genes SEPTIN9 and BCAR3.

## MATERIALS AND METHODS

### Data Preparation

#### Transcription and epigenetic profiling from Human Epigenome Atlas

We examined the association between the differential usage of exons and epigenetic marks using RNA-seq and ChIP-seq data for histone modifications from the Human Epigenome Atlas (release 9) [20]. This is a rich compendium of biological profiling experiments and is part of the Roadmap Epigenomics project. Data was downloaded from the ENCODE portal [21] (https://www.encodeproject.org/). Included were epigenomes with datasets for the histone marks H3K27ac, H3K27me3, H3K36me3, H3K4me1, H3K4me3, and H3K9me3. We omitted tissues lacking biological replicates, being flagged for poor quality controls, or that had unclear developmental origin. For the sake of homogeneity, only embryonic and adult samples were retrieved. 19 epigenomes including one cell line, seven *in vitro* differentiated cells, two primary cells and nine tissues passing the described filters were categorized by their potency, the life stage at their harvesting time, and the germ layer from which they arise. Supplemental table 1 lists the tissues/cells studied here.

#### Annotation of gene body and flank regions

The NCBI human reference genome GRCh38 was selected to annotate the gene components of interest. The GTF-formatted reference file was retrieved and flattened following the strategy of Anders et al [22]. Initially, we clustered genes that mapped to the same genomic region into one gene cluster to prevent mapping redundancy. Then, we sorted the group of exons belonging to the genes of one gene cluster and extracted the unique exons using the HTSeq package [23]. Meanwhile, genes were considered as one gene cluster only if they mapped to the same strand. If any two exons from different genes mapped to the same genomic region, they were rearranged and assigned to a new non-overlapping classification of exons that mapped to the same region (see suppl. figure 2). Instances of duplicated genes or genes spanning more than one genomic region were also considered. In fact, each duplicated copy was numbered sequentially. Promoters were defined as the regions between −2000 bp upstream of the transcriptional start site and 0 bp of a gene/gene cluster region. The intron/exon flanking regions were defined as the range between 200 bp upstream and downstream from an exon’s start/end site following existing studies [24]. Data annotation for differentially modified histone was performed using BEDTools [25]. Meanwhile, exons with differential usage were annotated by HTSeq as a part of the DEXSeq pipeline for DEU analysis [22, 23].

### Differential Analysis

#### Differential Exon Usage analysis

Here, we considered the transcript and exon abundance in the polyA-plus RNA-seq alignment files downloaded from ENCODE in BAM format. BAM files were sorted lexicographically and converted to SAM format via SAMTools [26]. Using HTSeq, we obtained the read counts for flattened exons in each sample from SAM files and used those as input for DEU analysis with the Bioconductor package DEXSeq [22] for all possible pairs of samples that can be composed from the 19 epigenomes. As an example, the exon usage (DEU) of the two genes SEPTIN9 and BCAR3 in two selected tissues is shown in the second line of Fig. 1A and 1B.

**Fig. 1.**
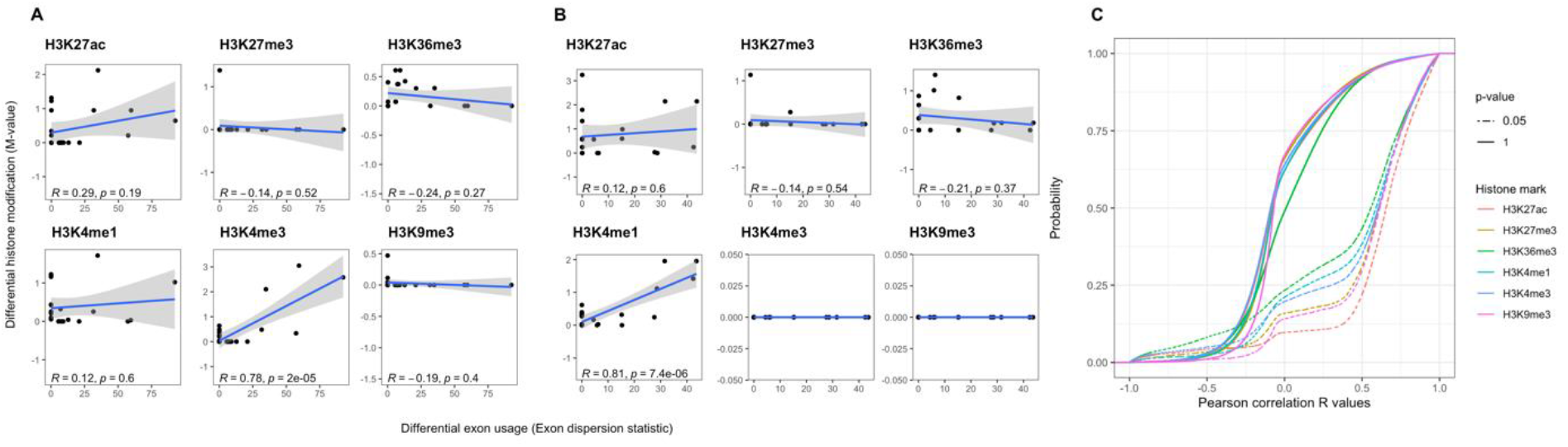
Pearson correlation between differential exon usage (detected by DEXSeq) and deregulation of six histone marks (absolute M-values detected by MAnorm) for (A) SEPTIN9, and (B) BCAR3. (C) shows the cumulative distribution of Pearson correlation for all genes. About 5 % of all genes show a correlation larger than 0.5 or smaller than −0.5. The dashed lines illustrate the cumulative distribution of all genes with top 5% highest correlation level (p=0.05).

#### Differential Histone Modification analysis

In order to account for putative technical noise in the data, we modelled the epigenomic read counts using regression analysis in a pairwise manner across all tissues. For this, we downloaded from ENCODE the GRCh38-assembled BAM-formatted alignments files and the BED-formatted replicated peaks or pseudo-replicated peaks files from Histone ChIP-seq analysis of H3K27ac, H3K27me3, H3K36me3, H3K4me1, H3K4me3, and H3K9me3 modifications, respectively. If multiple alignment files or peaks files existed for a specific histone type and epigenome, they were merged using SAMTools or BEDTools, respectively. All epigenomes were processed with the tool MAnorm for pairwise normalization and identification of differentially modified histone peaks [27]. MAnorm returns the log2 ratio of read density between two samples (M) and an adjusted p-value which we subsequently mapped to the flanking regions of each exon in the reference genome.

#### Combined differential expression analysis

Differential epigenetic profiles were then associated with exon rewiring by correlating the DEXSeq-generated DEU values for the entire gene to the respective absolute M-values from MAnorm mapped to the flanking regions around each exon boundary. To enhance the contrast between differential and non-differential features before correlation, all values for DEU and DHM with p-value larger than 0.05 were set to zero. Pearson correlation between differential features was computed for each gene between two tissues. Genes with correlation p-value below 0.05 are referred to as “epispliced genes” in our study. For illustration, Fig. 1 shows SEPTIN9 and BCAR3 as two well-known examples of epispliced genes, for which the relationship between AS and epigenetic modification was shown before [11, 28].

#### Tissue specific epispliced genes

One of our main interests was to identify epigenetic regulation mechanisms specific to certain tissues. Thus, we examined to which degree the identified epispliced genes are similar across 19 epigenomes and what is the overlap between them. For each tissue, we collected the set of epispliced genes by pooling the genes with Pearson correlation p-value below 0.05 from all the pairs containing the tissue of interest. Then, we computed a Tissue Specificity Index (TSI), which has been used in various contexts to identify tissue-specific transcripts, enhancers or tRNAs [29]). It is defined as:

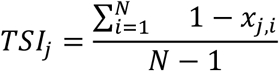

where, *N* is the number of tissues of interest and *x*_*j*,*i*_ is the RPKM in tissue *i* normalized over the maximum expression across the tissues for gene *j*. For each tissue and each epigenetic modification, Table 2 lists the number of epispliced genes with TSI larger than 0.75.

#### Association between epispliced genes and human development

We gathered all pairs/clusters of epigenomes that share the same epispliced genes/cluster genes for each histone modification type and report their number. All tissue-specific epispliced genes collected for each histone mark were subjected to GO term enrichment analysis of the biological process hierarchy using the PANTHER classification system [30] with a cutoff FDR-corrected p-value of 0.001. Enriched GO terms were sorted in decreasing order of fold enrichment and categorized into the four subgroups: cell responses and signaling, metabolic processes, developmental processes and other processes.

## RESULTS AND DISCUSSION

Differential placement of chromatin marks has been shown to substantially impact post-transcriptional processes including alternative splicing [16, 17, 31, 32]. With the aim of delineating their role in human development, we identified those genes where differential exon usage is correlated to the degree of histone mark deregulation at the exon boundaries. As exon boundaries we considered the flanking regions spanning 200 bp up-/downstream from the exon start and end points. We focused on the top 5% of such correlations. As examples, Fig. 1 shows the Pearson correlations for the genes SEPTIN9 and BCAR3. For SEPTIN9, only the mark H3K4me3 shows a significant correlation. For BCAR3, H3K4me1 shows a significant correlation. For comparison, Podlaha et al. reported Spearman rank correlations of histone mark levels and splicing exon inclusion rate of at most 0.36 for six normal human cell lines [33].

From now on, we will use the term “epispliced genes” to refer to genes showing correlations larger than 0.5. Table 1 lists all detected epispliced genes for the 19 tissues studied here. The largest numbers are found for the stem cells H1, mesenchymal stem cells and neuronal stem cells. The fewest events are found for the differentiated tissues aorta, small intestine, and sigmoid colon. Yet, all variations are within a factor of two. As a side note, we found that exon-body epigenetic effects appeared to be more pronounced than intronic or promoter effects. However, we did not analyze this in depth.

**Table 1.** Number of identified epispliced genes across 19 tissues/cells selected from the Human Epigenome Atlas for the six different histone marks listed at the top. Epispliced genes are those where differential exon usage overlaps with differential histone mark levels within +− 200 bp of the respective exon/intron boundaries and the correlation exceeds 0.5 or is below −0.5. (−) denotes cases where ChIP-seq histone peaks data were not available.

| Tissue | H3K27ac | H3K27me3 | H3K36me3 | H3K4me1 | H3K4me3 | H3K9me3 |
| --- | --- | --- | --- | --- | --- | --- |
| Adipose tissue | 1512 | - | - | - | - | - |
| Aorta | 1399 | 498 | 1089 | 1255 | 1367 | 380 |
| CD4-positive alpha beta T cell | 1931 | 897 | 1561 | 2056 | 1873 | 987 |
| CD8 positive alpha beta T cell | 2212 | 771 | 1696 | - | 2061 | 692 |
| Ectodermal cell | 2163 | 882 | - | - | 2159 | 696 |
| Endodermal cell | 2412 | 918 | 1681 | 2160 | 2334 | 970 |
| Esophagus | 1510 | 582 | 1075 | 1305 | 1554 | 505 |
| H1 cell | 2811 | 1464 | 1966 | 2486 | 2560 | 975 |
| Mesenchymal stem cell | 2551 | 947 | 1829 | 2039 | 2201 | 879 |
| Mesendoderm | 2717 | 896 | 2130 | 2387 | 2538 | - |
| Mesodermal cell | 2163 | - | 1625 | 1874 | 2174 | 778 |
| Neuronal stem cell | 2722 | 1275 | 1909 | 2514 | 2534 | 1475 |
| Pancreas | 1661 | 598 | 1531 | 1627 | 1558 | 673 |
| Psoas muscle | 2008 | 707 | 1343 | 1659 | 2038 | 501 |
| Sigmoid colon | 1604 | 606 | 1379 | 1658 | 1478 | 495 |
| Small intestine | 1441 | 596 | 1390 | 1619 | 1166 | 562 |
| Spleen | 1516 | 619 | 1419 | 1472 | 1375 | 729 |
| Stomach | 1531 | 734 | 1345 | 1481 | 1432 | 796 |
| Trophoblast | 2640 | 906 | 2031 | 2421 | - | 907 |

Fig. 2 illustrates the transcript architecture of the same two genes SEPTIN9 and BCAR3. Panels A and B were generated by the DEXSeq tool and illustrate overall counts (expression) in the top line, normalized counts for the replicate experiments in the third line, and the exon usage derived from this in the second line. Reassuringly, the two replicates show very similar normalized counts. The bottom line illustrates the genomic location of the exons. All cases of differential exon usage are marked in pink. For SEPTIN9, exons 4 and 11 are clear examples of strongly differential exon usage. For BCA3, this is most evident for exon 12. Panels C and D illustrate histone modification levels. All cases where differential exon usage overlaps with significant deregulation of histone marks are surrounded by boxes. For SEPTIN9 (C), the left-most box encloses exon 4. As discussed in Fig. 1 (A), the H3K4me3 mark is significantly correlated with DEU. Fig. 2 (C) shows this mark in the second-lowest row. One can clearly see that the H3K4me3 level in spleen (blue) is much higher than in neuronal stem cells (red).

**Figure 2.**
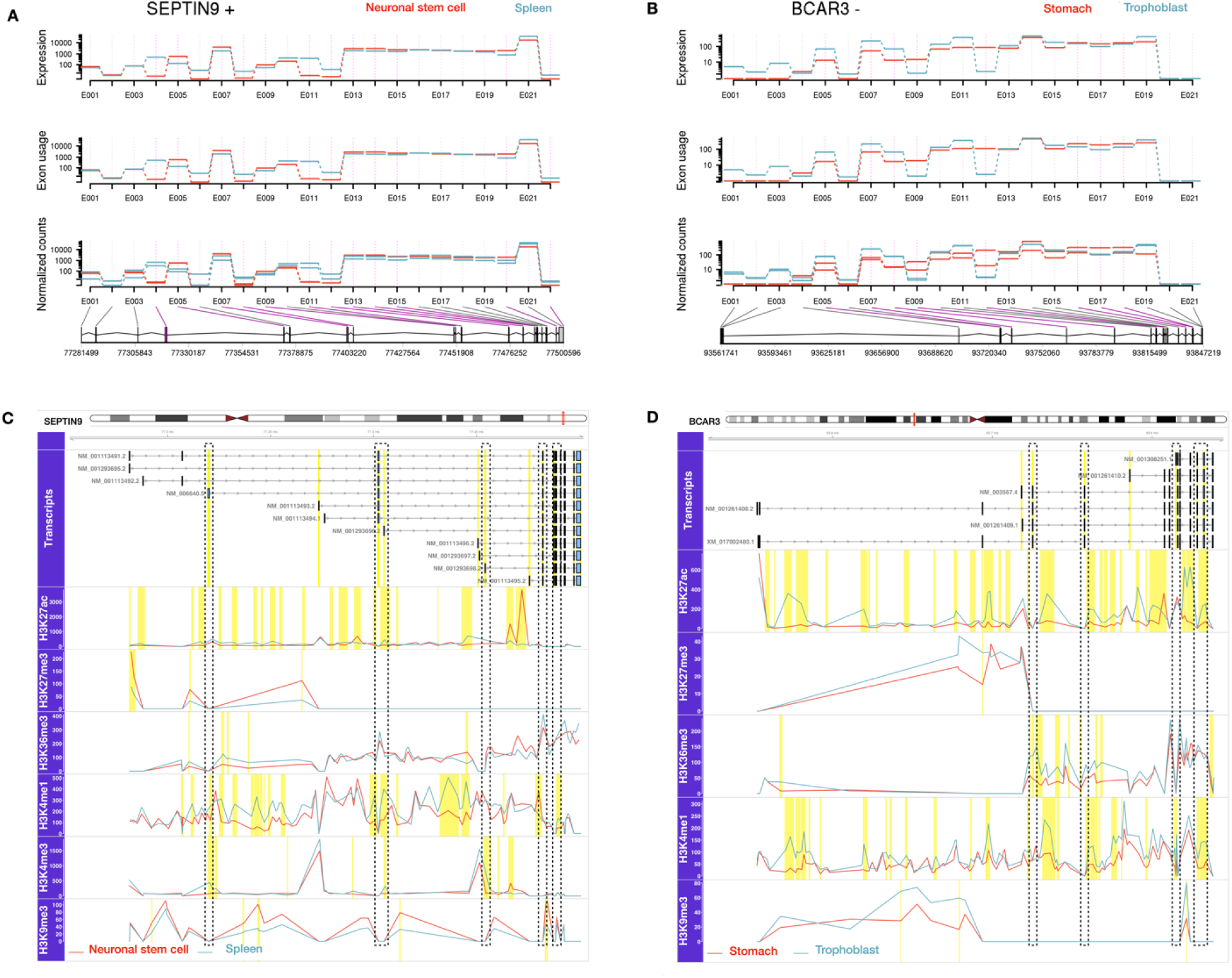
Two case studies (SEPTIN9 and BCAR3 genes) of epigenetic modifications associated with alternative splicing. (A) and (B) show diagrams produced by the DEXSeq package highlighting differential exon usage of the SEPTIN9 gene between neuronal stem cells and spleen and of the BCAR3 gene between stomach and trophoblast cells. Significantly differentially used exons (DEUs with FDR-adjusted p-value < 0.05) are marked in pink. (C) and (D) illustrate the association of DEUs and epigenomic modifications for the same two genes and tissues. Regions highlighted in yellow represent exons with differential usage identified by DEXSeq (FDR-adjusted p-value < 0.05) and significantly differentially enriched peaks for the histone modifications H3K36me3, H3K27ac, H3K27me3, H3K9me3, H3K4me3, and H3K4me1 detected by MAnorm (p-value < 0.05, |M-value| >= 1). The boxes show transcript variants found in the investigated cells or tissues as retrieved from NCBI Refseq. The figure was generated using the Gviz package.

The main aim of this paper was to analyze how episplicing and development are related to each other. Thus, we were less interested in detecting genes that are alternatively spliced in a similar manner in a large number of tissues. Rather, we wanted to focus on those genes showing differential exon usage coupled to epigenetic rewiring in a small set of tissues. Exactly such a bias is captured by the tissue-specific index (see methods). Table 2 shows the subsets of tissue-specific epispliced genes that were detected in a particular tissue. Obviously, these genes are subsets of Table 1. Sometimes, a gene may show correlated DEU and histone mark levels for multiple histone marks. The last column counts the total number of tissue-specific epispliced genes when overlaps are omitted. As in Table 1, the three stem cells show the largest number of tissue-specific epispliced genes, and aorta and sigmoid colon are among the ones having the fewest such genes (together with stomach and small intestine).

**Table 2.** Number of tissue-specific epispliced genes (TSI > 0.75) across all tissues in different epigenomics contexts. Genes with AS events correlated to differential modification of H3K27ac, H3K27me3, H3K36me3, H3K4me1, H3K4me3 and H3K9me3 were filtered by TSIs calculated from their normalized expression levels. The numbers of genes obtained for separate histone marks were summed together, either with and without inclusion of repeating cases to create the Total epispliced genes columns. (−) denotes cases where ChIP-seq histone peaks data were not available.

| Tissue | H3K27ac | H3K27me<br>3 | H3K36me<br>3 | H3K4me1 | H3K4me3 | H3K9me3 | Total epispliced<br>genes (with<br>overlaps) | Total epispliced<br>genes (without<br>overlaps) |
| --- | --- | --- | --- | --- | --- | --- | --- | --- |
| Adipose tissue | 388 | - | - | - | - | - | 388 | 388 |
| Aorta | 346 | 206 | 287 | 322 | 362 | 138 | 1661 | 1107 |
| CD4-positive alpha<br>beta T cell | 485 | 397 | 435 | 531 | 458 | 320 | 2626 | 1599 |
| CD8 positive alpha<br>beta T cell | 522 | 314 | 433 | - | 502 | 210 | 1981 | 1322 |
| Ectodermal cell | 596 | 408 | - | - | 616 | 224 | 1844 | 1237 |
| Endodermal cell | 637 | 404 | 484 | 573 | 598 | 279 | 2975 | 1764 |
| Esophagus | 460 | 269 | 299 | 388 | 457 | 189 | 2062 | 1308 |
| H1 cell | 718 | 561 | 594 | 685 | 743 | 322 | 3623 | 2074 |
| Mesenchymal stem<br>cell | 638 | 393 | 492 | 509 | 547 | 256 | 2835 | 1751 |
| Mesendoderm | 679 | 412 | 551 | 629 | 683 | - | 2954 | 1813 |
| Mesodermal cell | 534 | - | 472 | 492 | 529 | 219 | 2246 | 1445 |
| Neuronal stem cell | 694 | 569 | 586 | 661 | 718 | 435 | 3663 | 2115 |
| Pancreas | 464 | 270 | 388 | 454 | 453 | 217 | 2246 | 1440 |
| **Psoas muscle** | 509 | 305 | 354 | 428 | 549 | 176 | 2321 | 1411 |
| **Sigmoid colon** | 405 | 266 | 320 | 397 | 358 | 155 | 1901 | 1232 |
| **Small intestine** | 375 | 250 | 321 | 403 | 315 | 159 | 1823 | 1201 |
| **Spleen** | 481 | 283 | 423 | 422 | 425 | 264 | 2298 | 1482 |
| **Stomach** | 415 | 317 | 342 | 345 | 372 | 232 | 2023 | 1304 |
| **Trophoblast** | 695 | 407 | 533 | 642 | - | 274 | 2551 | 1743 |

Next, we asked what tissues share most or fewest tissue-specific epispliced genes. For this, we computed pairwise similarities between pairs of tissues by taking their Jaccard index based on the tissue-specific epispliced genes listed in Table 2. E.g., the largest overlap of 431 shared tissue-specific epispliced genes exists between H1 cells and mesendoderm, see panel E, while their union of tissue-specific epispliced genes is 995 genes. This then gives a Jaccard similarity of 0.433. As this value is not very high, there are clear differences in isoform expression between any two tissues. On the other hand, there are also clear similarities between certain tissue pairs. Thus, for each mark, we applied hierarchical clustering to group tissues with higher similarities into clusters.

We labeled the tissues by their potency, type, origin and life stage. To quantify which labeling type was associated most strongly with the clustering obtained, we computed adjusted Rand indices that quantify how well the labeling scheme matches the clustering results, see Table 3.

**Table 3.** Similarity between heatmap hierarchical clustering and tissue labelling schemes. The listed adjusted Rand indices measure how well the labelling of the investigated tissues according to their potency, sample type, origin, and life stage matches the hierarchical clustering according to the degree of shared tissue-specific epispliced genes. Precisely, the pairwise distance between any two tissues was calculated by subtracting their Jaccard similarity index from 100%. A Rand index of 1 would reflect a perfect clustering where all same-labeled tissues/cells are grouped in the same cluster, and differently labeled tissues/cells into different clusters. This analysis was done separately for the six histone marks investigated.

|  | Number of clusters considered | H3K27ac | H3K27me3 | H3K36me3 | H3K4me1 | H3K4me3 | H3K9me3 |
| --- | --- | --- | --- | --- | --- | --- | --- |
| Number of available tissues | - | 19 | 17 | 17 | 16 | 17 | 17 |
| Potency | 2 | 0,355 | 0,292 | 0,707 | 0,463 | 0,707 | 0,292 |
| Type | 3 | 0,519 | 0,49 | 0,381 | 0,473 | 0,381 | 0,75 |
| Origin | 3 | 0,218 | 0,27 | 0,475 | 0,342 | 0,366 | 0,27 |
| Life stage | 2 | 0,438 | 0,382 | 0,765 | 0,532 | 0,485 | 0,382 |

Fig. 3 shows a clustered heatmap of the similarity of tissue-specific epispliced genes between pairs of tissues. For H3K27me3 (panel B), H3K4me1 (D), and H3K9me3 (F), only relatively small similarities were found between all cells and tissues. For the H3K27ac mark (panel A), the largest similarities were found between neuronal stem cells, H1 cells, and mesendoderm as well as between C4 and CD8 immune cells. Differentiated tissues showed again low similarity among each other and with multipotent and pluripotent cells. According to the analysis for the H3K27ac mark in table 3 (panel A), samples having the same type shared most tissue-specific epispliced genes (Rand index 0,519). This is reflected by the fact that all differentiated tissues (except for psoas muscle) were clustered together. For H3K36me3 (C) and H3K4me3 (E), CD4 and CD8 cells shared high similarity as for H3K27ac. For these two marks, one also observes a cluster of six pluripotent and multipotent cells (neuronal stem cells, H1 cells, trophoblast or ectodermal cell, mesendoderm, mesodermal cell, endodermal cell) sharing fairly high similarity. This matches the Rand indices of table 3 that show high clustering purity for these two marks according to potency (0,707) as well as according to life stage for H3K36me3 (0,765).

**Figure 3.**
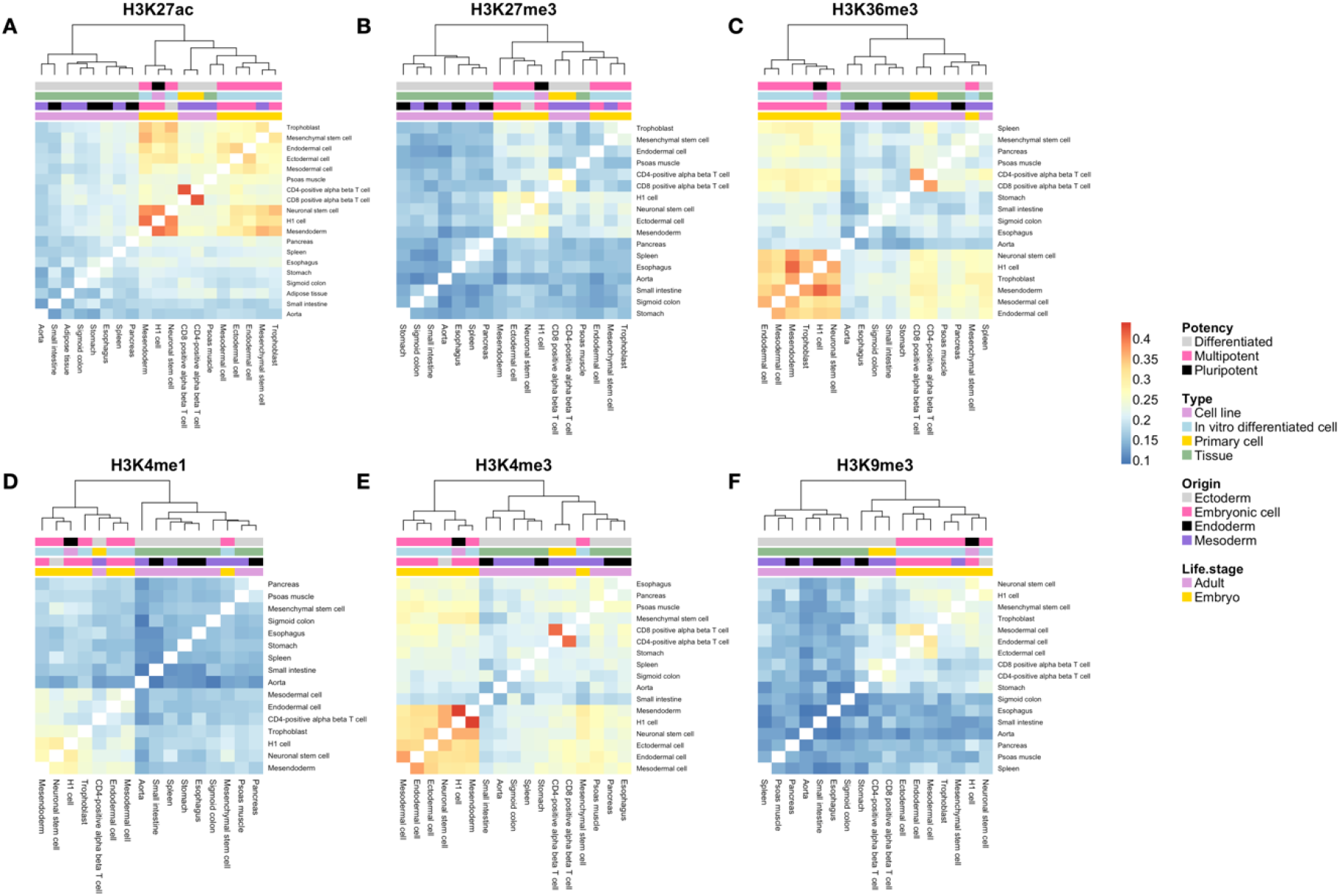
Heatmap representing hierarchical clustering of tissue-specific epispliced genes for the six histone marks H3K27ac, H3K27me3, H3K36me, H3K4me1, H3K4me3 and H3K9me3 (A-F). In each panel shows the pairwise similarities between 19 tissues. The similarities were measured by the Jaccard index, which is the ratio between the number of mutual genes and the total number of genes in the union sets of two tissues. All heatmaps use the same color scale ranging from 0.1 to the highest Jaccard index across all tissue pairs and histone modifications. Investigated tissues were annotated by their differentiation potency, type of sample, germ layer origin and the life stage when their samples were taken.

According to Fig. 3, stem cells and multipotent cells show a fairly high similarity of tissue-specific epispliced genes for the three marks H3K27ac, H3K36me3, and H3K4me3, whereas differentiated cells tend to have rather low similarities overall. The only exception to this are CD4 and CD8 cells that also have high similarities for the same three histone marks, but not for the other three marks.

Finally, we performed functional enrichment analysis of the tissue-specific epispliced genes separately for each histone mark. Fig. 4 shows the results of gene-set enrichment analysis based on the biological process branch of the Gene Ontology. The terms are arranged into four broad GO-SLIM categories, namely cell response and signaling, metabolic processes, developmental processes, and other processes. The category of developmental processes plays a dominant role. The two marks H3K27me3 and H3K4me3 seem to have the largest contribution in this. Interestingly, these two marks showed very different behavior in terms of tissue-similarity in Fig. 3. Whereas the tissues and cells shared little similarity for the H3K27me3 mark in terms of shared tissue-specific epispliced genes (Fig. 3), they had noticeably higher similarities for the H3K4me3 mark. Epigenetic regulation of differential exon usage is also important for several rather general metabolic processes, but only for the three marks H3K27ac, H3K36me3, and H3K4me1, which are disjoint from the two marks (H3K27me3 and H3K4me3) that mostly control developmental processes. The H3K9me3 mark is set apart and seems to play a specific role in regulating cell response and signaling processes.

**Figure 4.**
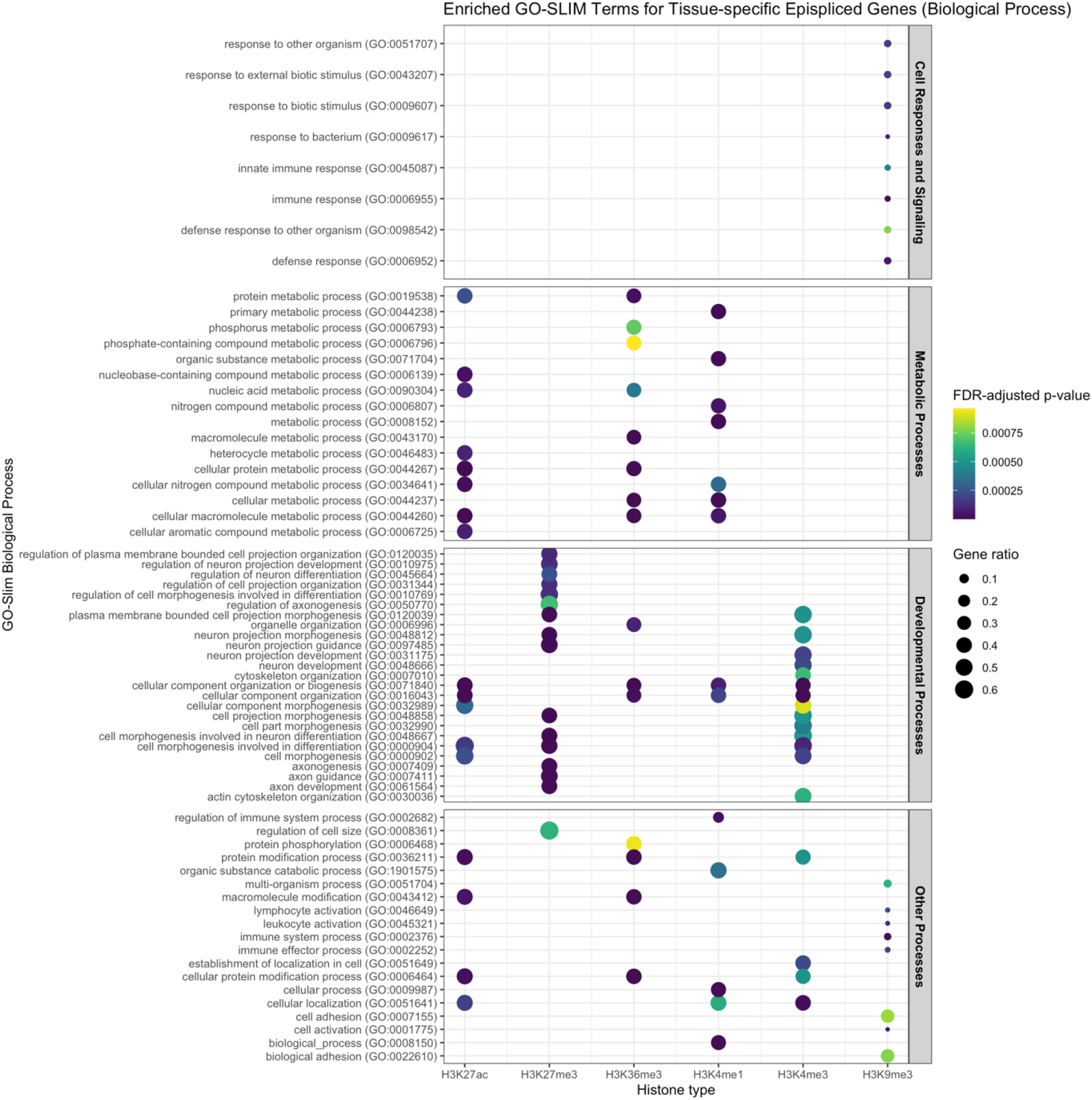
Gene ontology (GO) enrichment analysis for biological functions of tissue-specific epispliced genes for the six histone marks. In each case, the top enriched GO terms (FDR-adjusted p-value < 0.001) were sorted in decreasing order of significance and with respect to overlap between the histone marks.

For comparison, a previous study [39] found that genes with H3K27ac-enhanced regions are associated with GO functions that are characteristic for multipotent stem cells, such as anatomical structure development and nervous system development. Broad H3K4me3 domains were also reported with distinctive roles in neuron development during stem cells and human brain tissue differentiation which is in concordance with our findings [40]. Furthermore, H3K4me3 and H3K27me3 promoter bivalency was established as a prominent epigenetic mechanism for lineage-specific activation or repression of developmental genes in embryonic and neural stem cell differentiation [41, 42]. Meanwhile, the GO terms for H3K4me1 cover a wider span of biological functions and contribute mainly to biogenesis and transcriptional activation. This agrees with the findings of Stunnenberg et al. who reported that pre-marking for enhancers activation succeeded mainly with H3K4me1 enrichment during stem cell differentiation [43]. For H3K36me3, many GO terms contribute to cellular component organization or RNA processing and regulation besides morphogenesis. This opens up the possibility that H3K36me3 contributes to developmental processes via transcriptional regulation. In mouse embryonic stem cells, crosstalk between H3K36me3 and the RNA modification m6A mediates the maintenance of pluripotency and the initiation of differentiation via recruitment of RNA methyltransferase complexes [44].

In summary, our findings emphasize the notion that exon-intron boundaries set by histones/epigenetic marks are not only used to define the ends of the elements for the mRNA transcript to be expressed. Rather, they are also part of a machinery that regulates and controls the relative abundance of different transcripts or protein isoforms that map to the same chromosomal region across tissues. We found that the coupling between histone marks and differential exon usage is most prominent in semi-early (embryonic and fetal) developmental stages and most frequently affects developmental genes. We are certain that our findings on the triangle alternative splicing – epigenetic deregulation – development will attract further attention in the near future. One important further element that we omitted from our analysis is to address the specific roles of individual splicing factors in this.

## SUPPLEMENTS

**Figure S1.**
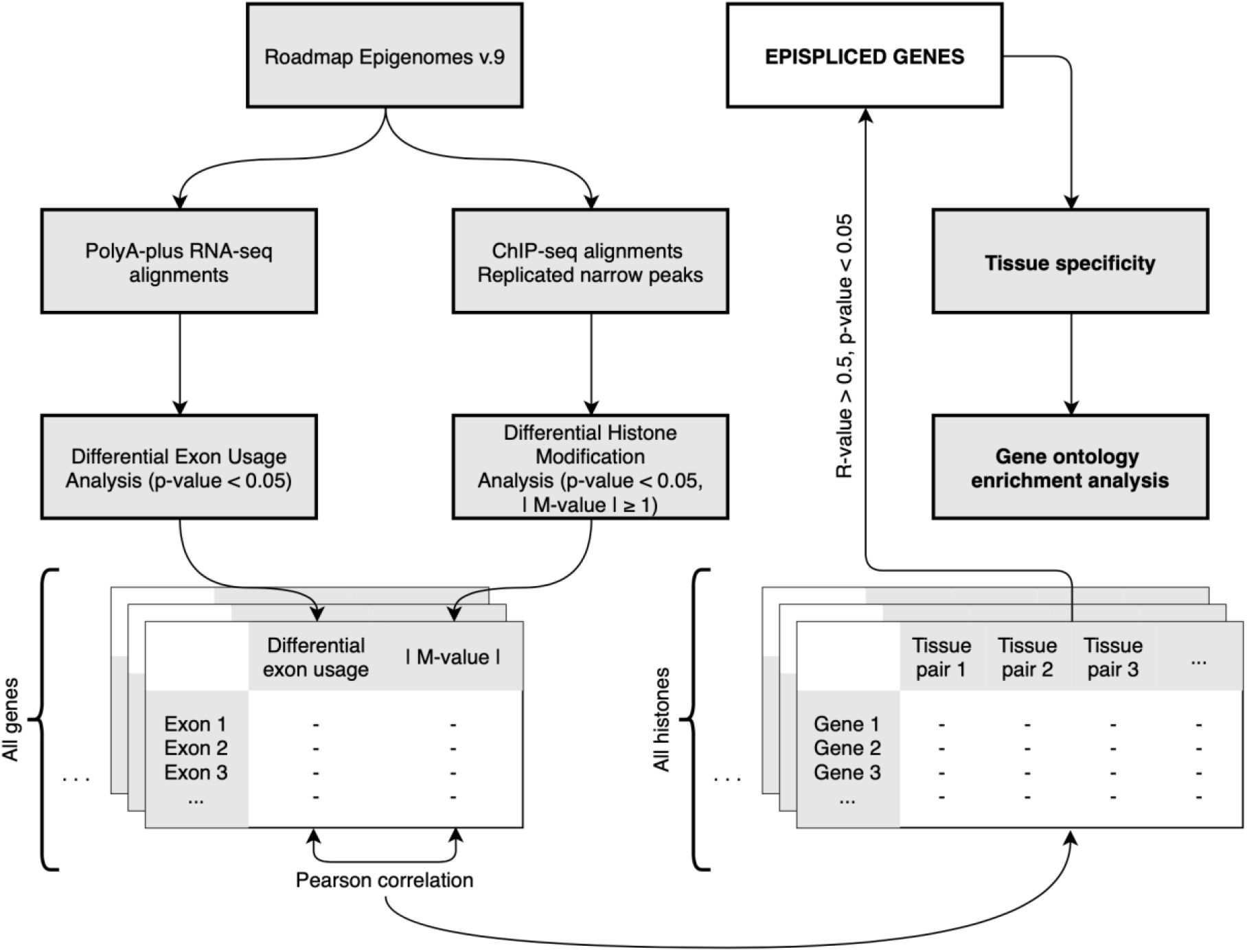
Schematic for workflow of the epispliced genes analysis. Expression data and histone enrichment data were collected from Roadmap Epigenomes project version 9 and were subjected to Differential Exon Usage and Differential Histone Modification analysis respectively. For each gene, the values from the two features were correlated by Pearson correlation. Epispliced genes are defined with R-value larger than 0.5 and FDR-adjusted p-value smaller than 0.05. Tissue specificity index was calculated for the epispliced genes. Gene ontology terms for biological processes enrichment analysis was performed for the tissue-specific epispliced genes collection of each histone.

**Figure S2.**
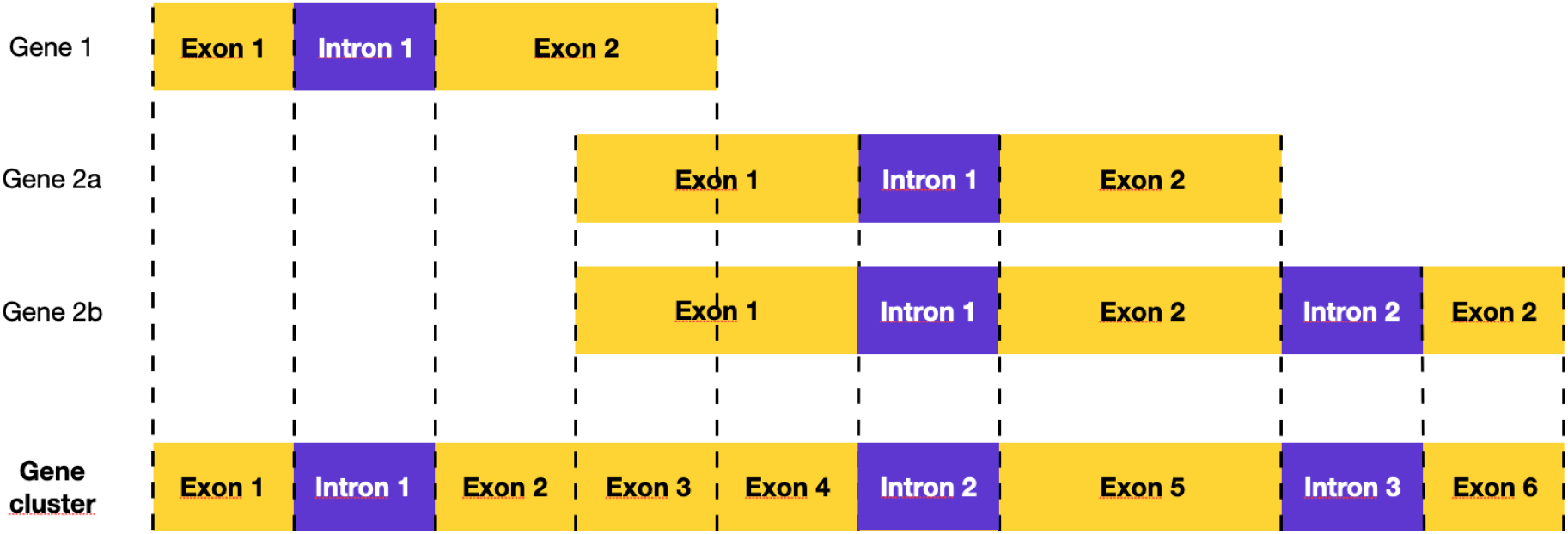
Flattening of overlapping genes and transcripts following DEXSeq scheme. Overlapping exonic regions are counted as a new exon in the collapsed gene cluster. Intronic regions are retained in the same manner. Numbering of exons and introns is according to their positions in the flattened gene cluster.

**Table S1.** List of tissue and cell types retrieved from Roadmap Epigenomes project with annotated potency, sample type, origin, and life stage.

| Epigenome | Potency | Type | Origin | Life stage |
| --- | --- | --- | --- | --- |
| Adipose tissue | Differentiated | Tissue | Mesoderm | Adult |
| Aorta | Differentiated | Tissue | Mesoderm | Adult |
| CD4-positive alpha beta T cell | Differentiated | Primary cell | Mesoderm | Adult |
| CD8-positive alpha beta T cell | Differentiated | Primary cell | Mesoderm | Adult |
| Ectodermal cell | Multipotent | In vitro differentiated cell | Embryonic cell | Embryo |
| Endodermal cell | Multipotent | In vitro differentiated cell | Embryonic cell | Embryo |
| Esophagus | Differentiated | Tissue | Endoderm | Adult |
| H1 cell | Pluripotent | Cell line | Embryonic cell | Embryo |
| Mesenchymal stem cell | Multipotent | In vitro differentiated cell | Mesoderm | Embryo |
| Mesendoderm | Multipotent | In vitro differentiated cell | Embryonic cell | Embryo |
| Mesodermal cell | Multipotent | In vitro differentiated cell | Embryonic cell | Embryo |
| Neuronalstem cell | Multipotent | In vitro differentiated cell | Ectoderm | Embryo |
| Pancreas | Differentiated | Tissue | Endoderm | Adult |
| Psoas muscle | Differentiated | Tissue | Mesoderm | Adult |
| Sigmoid colon | Differentiated | Tissue | Mesoderm | Adult |
| Small intestine | Differentiated | Tissue | Endoderm | Adult |
| Spleen | Differentiated | Tissue | Mesoderm | Adult |
| Stomach | Differentiated | Tissue | Endoderm | Adult |
| Trophoblast cell | Multipotent | In vitro differentiated cell | Embryonic cell | Embryo |

